# scCancer: a package for automated processing of single cell RNA-seq data in cancer

**DOI:** 10.1101/800490

**Authors:** Wenbo Guo, Dongfang Wang, Shicheng Wang, Yiran Shan, Jin Gu

## Abstract

**Summary:** Molecular heterogeneities bring great challenges for cancer diagnosis and treatment. Recent advance in single cell RNA-sequencing (scRNA-seq) technology make it possible to study cancer transcriptomic heterogeneities at single cell level. Here, we develop an R package named scCancer which focuses on processing and analyzing scRNA-seq data for cancer research. Except basic data processing steps, this package takes several special considerations for cancer-specific features. Firstly, the package introduced comprehensive quality control metrics. Secondly, it used a data-driven machine learning algorithm to accurately identify major cancer microenvironment cell populations. Thirdly, it estimated a malignancy score to classify malignant (cancerous) and non-malignant cells. Then, it analyzed intra-tumor heterogeneities by key cellular phenotypes (such as cell cycle and stemness) and gene signatures. Finally, a user-friendly graphic report was generated for all the analyses.

**Availability:** http://lifeome.net/software/sccancer/.

## 1 Introduction

Cancer is a kind of highly heterogeneous diseases. Cells from the same patient’s tumor may have different expression profiles. Recently, various single cell RNA-sequencing (scRNA-seq) techniques have been widely applied to study cancer heterogeneities at single cell level. Among these techniques, droplet-based platforms can profile thousands of cells at a time and are more appropriate for highly heterogeneous application scenarios (Zheng *et al.*, 2017).

Currently, many tools and algorithms have been developed to analyze scRNA-seq expression data. For example, Seurat is one of the most popular R packages and contains some basic analyses (Butler *et al.*, 2018). However, cancer samples have their own features, such as complex microenvironment and high intra-tumor heterogeneity. So, it is very useful to develop cancer-specific tools beyond the basic analyses.

Here, we developed a user-friendly and automated R package scCancer for cancer scRNA-seq data analysis. In the package, we encapsulated basic analyses and included more comprehensive quality control (QC) metrics. Besides, it integrated several specific computational analyses for cancer data: major cell type classifications of cancer microenvironment; cell malignancy estimation; intra-tumor heterogeneities for important cellular phenotypes (cell cycle and stemness); and gene signatures based heterogeneity analyses.

## 2 Methods

### 2.1 Workflow overview

The workflow of scCancer mainly consists of two parts. The first, named *scStatistics*, performed basic statistical analysis of raw data and quality control. The second, named *scAnnotation*, performed functional data analyses and visualizations, such as low dimensional representation, clustering, cell type classification, malignancy estimation, cellular phenotype scoring, gene signature analysis, etc. (Fig. 1).

**Fig. 1.**
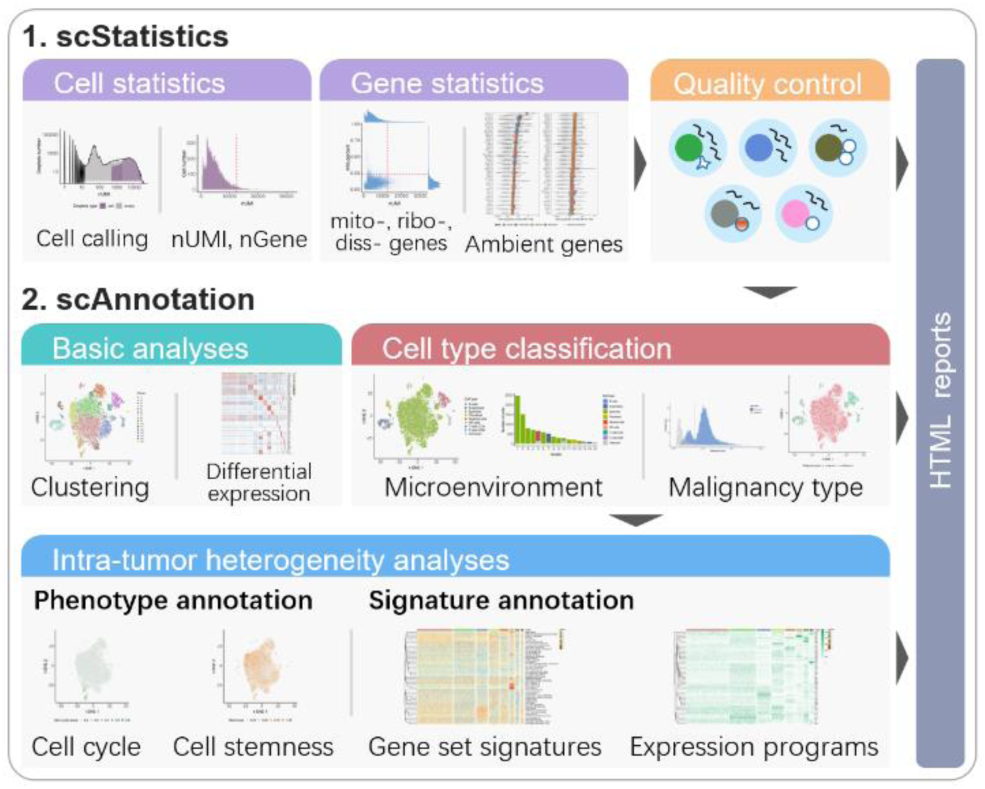
scCancer workflow.

### 2.2 Quality control

After cell calling, the droplets containing cells have been identified. However, droplets with low quality cells and more than one cell are still need to be exclude. Except some commonly used metrics, we introduced another metric to remove cells affected by solid tissue dissociation process (van den Brink *et al.*, 2017). Besides, considering that different sample sources and experimental conditions may lead to different distribution of the QC metrics, we adaptively determined the filtering thresholds by detecting outliers from the distribution of metrics, instead of routine fixed cutoffs (Supplementary Section S2).

For QC of genes, we found mitochondrial genes, ribosomal genes, and some other gene (such as MALAT1, FTH1, B2M) expressed high proportion in both cells and background in nearly all samples. These genes may significantly contribute to ambient RNAs. Here, we proposed to remove them before normalization (Supplementary Section S3).

### 2.3 Basic downstream analyses

After filtering, we performed downstream analyses based on the R package Seurat and redesigned the generated graphics. These analyses mainly include normalization, log-transformation, highly variable genes identification, removing unwanted variance, scaling, centering, dimension reduction, clustering, and differential expression analysis (Stuart *et al.*, 2019) (Supplementary Section S4).

### 2.4 Microenvironment cell type classification

Cancer microenvironment plays an important role in the tumor progression. Here, we developed a data-driven method to annotate major micro-environment cell types, including endothelial cells, fibroblast, and immune cells (CD4+ T cells, CD8+ T cells, B cells, nature killer cells, and myeloid cells). We curated a high-quality dataset by combining multiple cancer scRNA-seq data and trained one-class logistic regression (OCLR) machine learning models for different cell types (Sokolov *et al*., 2016). Based on the trained cell type templates, spearman correlations were used to classify different cell types (Supplementary Section S5).

### 2.5 Cell malignancy estimation

Estimating cell malignancy and distinguishing malignant and non-malignant cells is also a critical issue. Generally, copy number alterations (CNV) inferred from scRNA-seq data are potential to identify malignant tumor cells. Here, we first used the algorithm of R package infercnv (Patel *et al.*, 2014) to get an initial estimation of CNVs. Then, we took advantage of cells’ neighborhoods information to smooth CNV values and defined the malignancy score as the mean of the squares of them. By comparing the distribution of malignancy scores with reference and its bimodality, the malignant cells were identified (Supplementary Section S6).

### 2.6 Intra-tumor cell phenotype heterogeneity analyses

scCancer focused on two cellular phenotypes, cell cycle and stemness, to analyze intra-tumor heterogeneity. For cell cycle, the relative average expression of a list of G2/M and S phase markers was defined as cell cycle score (Stuart *et al.*, 2019). For cell stemness, a stemness signature was identified based on a stem/progenitor cells dataset using OCLR model. The stemness score was defined as the Spearman correlation coefficient between the signature and cells’ expression (Malta *et al.*, 2018) (Supplementary Section S8, S9).

### 2.7 Intra-tumor cell signature heterogeneity analyses

Gene signature analysis is commonly used to analyze tumor heterogeneities. scCancer used gene set variation analysis (GSVA) (Hänzelmann *et al*, 2013) for known gene set based signature analysis. An alternative method based on the relative average expression level across gene set was also provided. By default, scCancer used 50 hallmark gene sets from MSigDB and users can also input their own sets (Supplementary Section S10).

Besides, scCancer can also unsupervised identify potential expression program signatures. It applied non-negative matrix factorization (NMF) to the centralized and non-negative changed expression matrix. According to the decomposed matrixes, it can find potential expression programs and the cell sub-populations (Supplementary Section S11).

## 3 Discussion

Our R package scCancer allowed users to automatically analyze droplet-based cancer scRNA-seq data. It integrated basic single cell processes and cancer-specific analyses. In the future, we will provide more optional methods for each step and try to integrate more cancer-related functions.

