## Supplementary for "scCancer: a package for automated processing of single cell RNA-seq data in cancer"

### S1. Cell calling

The input of scCancer is a feature-barcode expression matrix generated by cell ranger. In addition to reads alignments and matrix generation, cell ranger v3 also uses EmptyDrops algorithm to perform cell calling [1]. When the numbers of RNA in cells are not balanced, droplets with low RNA content may be neglected. In order to identify these potential cells, EmptyDrops estimates a background distribution from nearly empty droplets (total UMI count $\leq 100$), and uses hypothesis test to distinguish more cells from ambiguous droplets. Comparing cell ranger v2, it is more appropriate for tumor samples which contain large-size tumor cells and small-size immune cells. So, we suggested users used cell ranger v3 to generate raw matrix. Even so, we also provided corresponding process to achieve compatible with cell ranger v2.

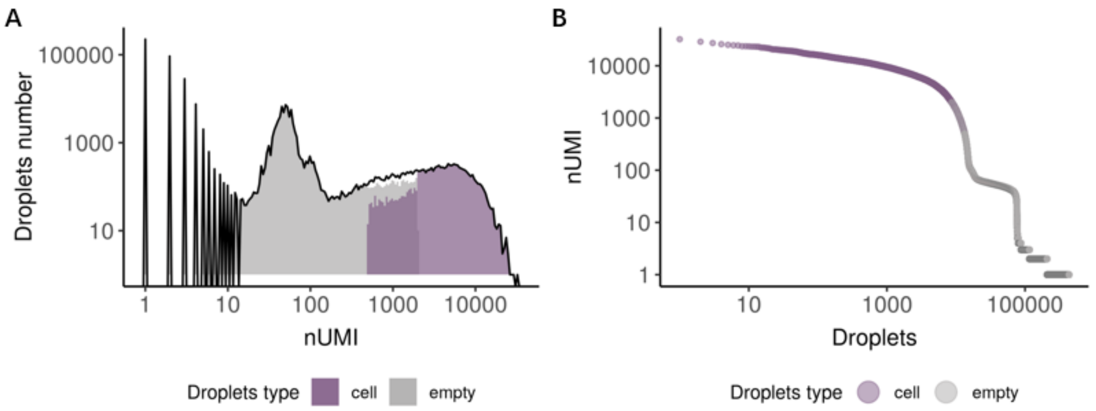


**Figure S1**. The visualization results of cell calling in scCancer.
**(A)** The histogram for the distribution of total UMI count in putative cells (purple) and empty droplets (grey).
**(B)** The rank plot for the droplets' total UMI count (nUMI).

### S2. Cell quality control

Ideally, one droplet contains one cell in good state and the detected RNA transcripts are all from this cell. However, some abnormal situations may occur during the single cell isolation procession (Fig. 1).

- The droplet contains one cell but it is contaminated by some ambient RNAs from necrotic or lysed cells so that the expression profile is influenced.
- The droplet contains a necrotic or lysed cell and many nuclear RNA transcripts are lost so that the percentage of transcripts from mitochondrial genes is relatively high.
- The droplet doesn't capture a cell and only capture some ambient RNAs so that the total UMIs in the droplet is very small.
- The cells are not isolated completely, and one droplet contains two or more cells (doublets or multiplets) so that the total UMIs in the droplet is extremely large.
- Because solid tumor samples are usually surrounded by connective tissue, it is easy to induce some cells to produce stress response during sample dissociation process so that cells may highly express some genes, such as immediate early genes and heat-shock proteins genes [2]. So, if cells are affected seriously, the dissociation-associated genes may

have a large percent in the total UMIs. In scCancer, we referred to the results of van den Brink, and defined a dissociation-associated gene list [2].

Considering these abnormal situations, we calculated following metrics to perform quality control (QC).

- nUMI: the number of total UMIs in the droplet. Too small means no cells are captured, and too large means capturing two or more.
- nGene: the number of expressed genes in the droplet. Too small means the loss of transcripts diversity. Too large means containing two or more cells.
- mito.percent: the percent of UMIs from mitochondrial genes. Too large means the captured cell is necrotic or lysed.
- ribo.percent: the percent of UMIs from ribosome genes. Too large means the captured cell is necrotic or lysed.
- diss.percent: the percent of UMIs from dissociation-associated genes. Too large means dissociation process has a serious effect on the cell.

By testing on some different samples, we found the distributions of above metrics were not very similar across samples from different sources and experimental environment. So, we proposed to determine the filter upper thresholds according to the distribution instead of routine fixed value. Specifically, we used the function “boxplot.stats” of R package “grDevices” to detect outliers and set the minimum of outliers as upper thresholds. Besides, the lower thresholds of nUMI and nGene, we set 3 and 200, respectively.

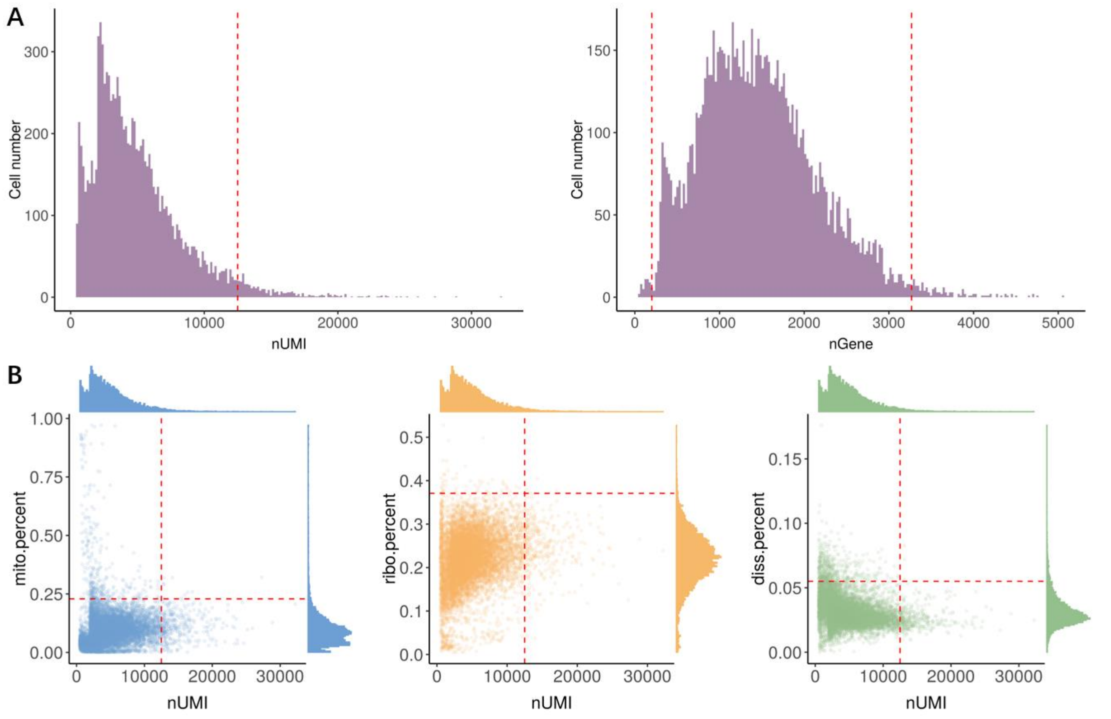


**Figure S2**. The visualization results of cell QC in scCancer. The red dash line marks the identified thresholds for QC.

**(A)** The histograms for the distribution of nUMI (left) and nGene (right).

**(B)** A scatter plot shows the relationship between three types of genes’ percentage and nUMI. The marginal plots are the corresponding histograms for its axes. The left one is for mitochondrial genes (blue). The middle one is for ribosome genes (yellow). The right one is for dissociation-associated genes (green).

### S3. Gene quality control

As mentioned in cell quality control, some necrotic or lysed cells may leak its RNA transcripts into the external suspensions, so that other droplets may be contaminated by these ambient RNA transcripts. In order to analyze the influence of contamination, we performed statistics on several samples. Firstly, we constructed a background distribution. We assumed that the droplets with nUMI less than $T$ (default 10) were empty and their UMIs were all from ambient RNAs. By summing the number of UMIs across these empty droplets, we got a background distribution and the proportion of each gene can be treated as its probability of being contaminated. Secondly, we calculated the proportion of UMIs for each gene in each cell. And we got a figure like Fig. S3A, which showed the top 100 genes' proportions distribution in cells by boxplot (ordered by median) and the red star signs marked genes' proportion in background. Besides, we can also find that it is extremely correlated for background proportion, median of proportion in cells, and gene detection rate in cells (Fig. S3B and S3C). By testing on some different samples, we got similar results that mitochondrial genes (blue), ribosomal genes (yellow), dissociation-associated genes (green), and some other genes (such as MALAT1, FTH1, B2M, and so on, red) were always ranked steadily in the forefront of all genes. We performed functional enrichment analysis on these genes (red), we found they were mainly related to membrane and ATP synthase complex (Fig. S3D). So, we can get conclusions that these genes have large proportion in background and are more likely to be imported into droplets from outside. At the same time, they perform basic functions within cells and have less influence on downstream analysis. Therefore, we suggested to remove these genes before normalization in the scCancer pipeline.

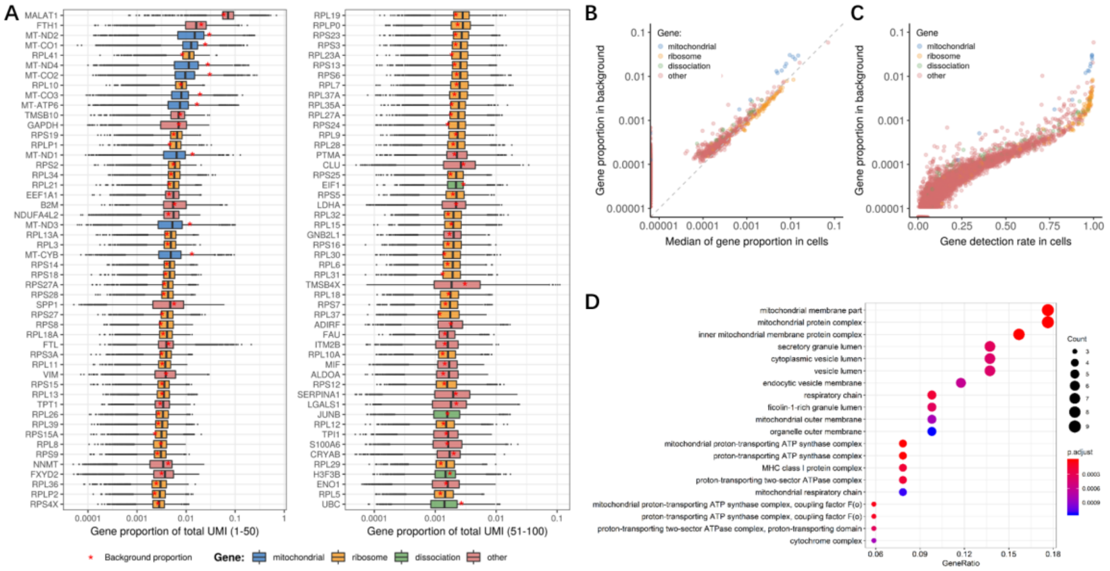


**Figure S3**. The visualization results of gene QC.

**(A)** A boxplot shows the distributions of some genes' expression profiles. The x axis is the gene proportion of total UMI for cells. The y axis is the top 100 genes ordered by the median. In each box, red star signs mark the genes' proportion in background. The color of boxes means four type of genes.

**(B)** A scatter plot shows the relationship between gene proportion in background and the median of gene proportion in cells.

**(C)** A scatter plot shows the relationship between gene proportion in background and the gene detection rate in cells.

**(D)** The results of functional enrichment analysis for the genes with high background proportion (genes with red color in A, B, C) in different samples.

### S4. Basic downstream analyses

After QC, we performed some basic downstream analyses using R package Seurat [3], which mainly included normalization, log-transformation, highly variable genes (HVG) identification, removing unwanted source of variance, scaling, centering, dimension reduction, clustering, and differential expression analysis. Here, we set the arguments of Seurat functions "pc.use", "resolution" as 30 and 0.8, respectively. Besides, we also redesigned the generated graphics to make them be able to automatically adjust according to the characteristics of data.

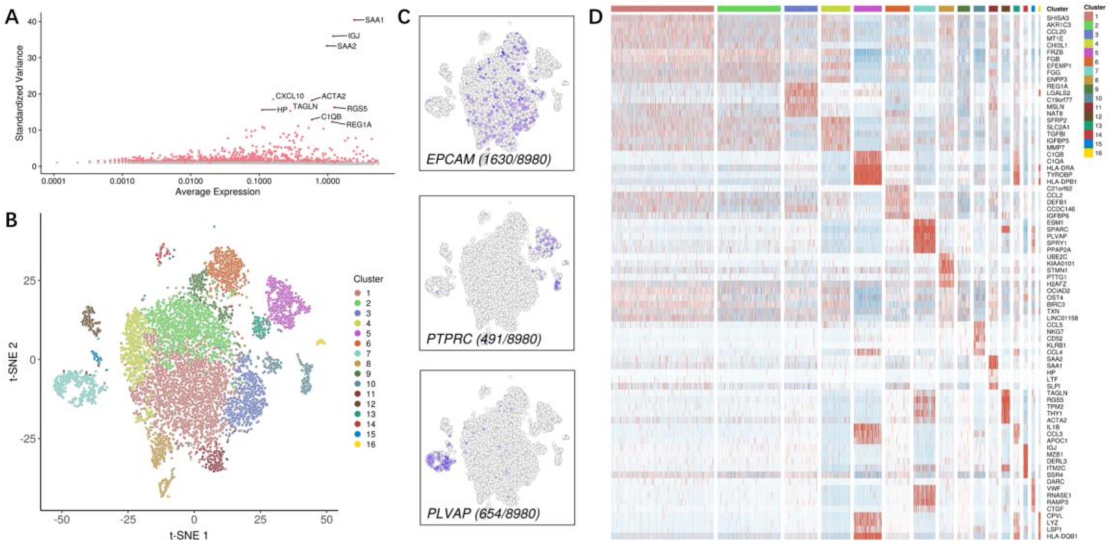


**Figure S4**. The visualization results of routine downstream analyses in scCancer.

**(A)** A scatter plot shows the highly variable genes.

**(B)** A t-SNE plot shows the clustering results.

**(C)** Some t-SNE plots show the expression profile of three representative genes. The bottom left label is the gene's name and its detection rate.

**(D)** A heatmap shows the expression profile for the top 5 differentially expressed genes for every cluster compared to all remaining cells. The color of top bar represents different clusters.

### S5. Cancer microenvironmental cell type classification

In order to annotate cancer microenvironmental cell types, we proposed a data-driven one-class logistic regression (OCLR) machine learning model [4], which learned from a single class training set and was appropriate to the situation absenting negative samples.

We selected some common cell types in solid tumor, epithelial cells, endothelial cells, fibroblast, and immune cells (CD4+ T cells, CD8+ T cells, B cells, nature killer cells, and myeloid cells), and collected some single cell samples which had been labeled with these types. The number of training cells for these types is presented in the Table S1.

**Table S1.** The number of training cells for each type.

| Cell types | The number of training cells |
|---|---|
| Epithelial cells | 9204 |
| Endothelial cells | 792 |
| Fibroblast | 264 |
| CD4+ T cells | 1613 |
| CD8+ T cells | 1919 |
| B cells | 425 |

| | |
|---|---|
| Nature killer cells | 308 |
| Myeloid cells | 1066 |

For each cell type, we trained an OCLR model on the corresponding training cells respectively. Formally, we set the above eight sample sets as $\mathbf{X}^{(k)} \in R_{m_k * n_k}$, $k = 1, 2, \ldots, 8$, where $m_k$ is the number of shared genes in set $k$, and $n_k$ is the number of samples in set $k$. Then the aim of OCLR is training the weight vector $\boldsymbol{w}_{(k)} \in R_{m_k * 1}$ to maximize the log-likelihood:

$$l\left(w_{(k)} \mid \mathbf{X}^{(k)}\right) = \Sigma_{i=1}^{n_k} \log p\left(\boldsymbol{x}_i^{(k)} | \boldsymbol{w}_{(k)}\right) ,$$

where $p\left(\boldsymbol{x}_i^{(k)} | \boldsymbol{w}_{(k)}\right)$=$\frac{\exp\left(\boldsymbol{w}_{(k)}^T \boldsymbol{x}_i^{(k)}\right)}{1+\exp\left(\boldsymbol{w}_{(k)}^T \boldsymbol{x}_i^{(k)}\right)}$. In order to avoid $\boldsymbol{w}_{(k)}$ tends to infinity, OCLR imposes a regularizer $R\left(\boldsymbol{w}_{(k)}\right)$ . Here, we set $R\left(\boldsymbol{w}_{(k)}\right) = \frac{1}{2}\left(\boldsymbol{w}_{(k)}^T \boldsymbol{w}_{(k)}\right)$. So, the final objective function is:

$$\frac{1}{n} \Sigma_{i=1}^{n_k} \left(\boldsymbol{w}_{(k)}^T \boldsymbol{x}_i^{(k)} - \log\left(1 + \exp\left(\boldsymbol{w}_{(k)}^T \boldsymbol{x}_i^{(k)}\right)\right)\right) - \frac{1}{2}\left(\boldsymbol{w}_{(k)}^T \boldsymbol{w}_{(k)}\right).$$

The training process can be performed by coordinate descent. In scCancer, we used R package gelnet to solve it [5]. Using the trained eight gene weight vectors $\boldsymbol{w}_{(k)}$, we assigned eight cell type scores to each cell by calculating the Spearman correlation coefficients between $\boldsymbol{w}_{(k)}$ and the cell's expression vector. If one of the eight scores were significantly larger than others, we annotated the cell with the corresponding type. If the eight scores were all less than 0.1 or were all located in the first 0.1 quantile, we annotated with "Unknown".

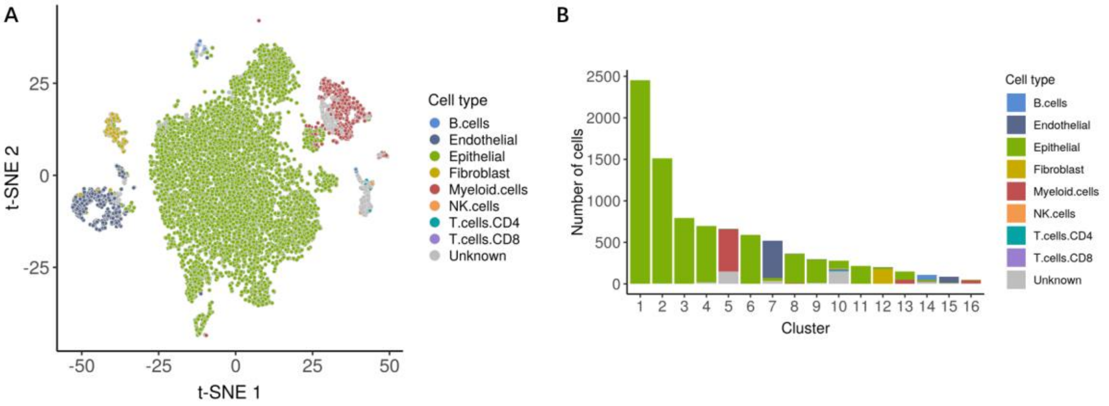


**Figure S5**. The visualization results of cancer microenvironmental cell annotation in scCancer.
**(A)** A t-SNE plot shows the predicted cell types.
**(B)** A bar plot shows the proportion of different cell types in each cluster.

### S6. Cell malignancy estimation

Here, we referred to the CNV inference algorithm of R package infercnv and used the CNV profile to measure cell malignancy degree [6].

In order to estimate CNV, we prepared a default hg19 gene position information file, and a reference normal cells set (an expression matrix with 2552 cells, named as reference) in scCancer. For the test sample matrix (named as observation), firstly, we removed genes with low mean expression or small detection rate. Secondly, we performed normalization, Anscombe transform, log-transform on it and set lower and upper bounds to limit outliers. Then we did convolution on each chromosome using a window of 101 genes. After centering across chromosomes by median, subtracting the reference profiles, inverting log-transformation, and removing outliers, we got an initial estimated CNV profile matrix.

Considering the influence of dropout, we proposed to use cells' neighborhood relationship to make the CNV profile more complete. In detail, we took advantage of the similarity matrix of clustering to identify the neighborhoods of each cell, and calculated the weighted average as the final CNV profile. Then we defined the malignancy score as the mean of the squares of weighted CNV values across chromosome.

Besides, in order to classify cells into the malignant and non-malignant, we proposed an algorithm to detect the bimodality of malignancy scores distribution. Firstly, if the scores of sample observation and normal cells reference shared a common single peak, we assigned non-malignant labels to all cells. Otherwise, if the distribution is bimodality (like Fig S6A), we detected the largest valley to classify cells into non-malignant and malignant clusters.

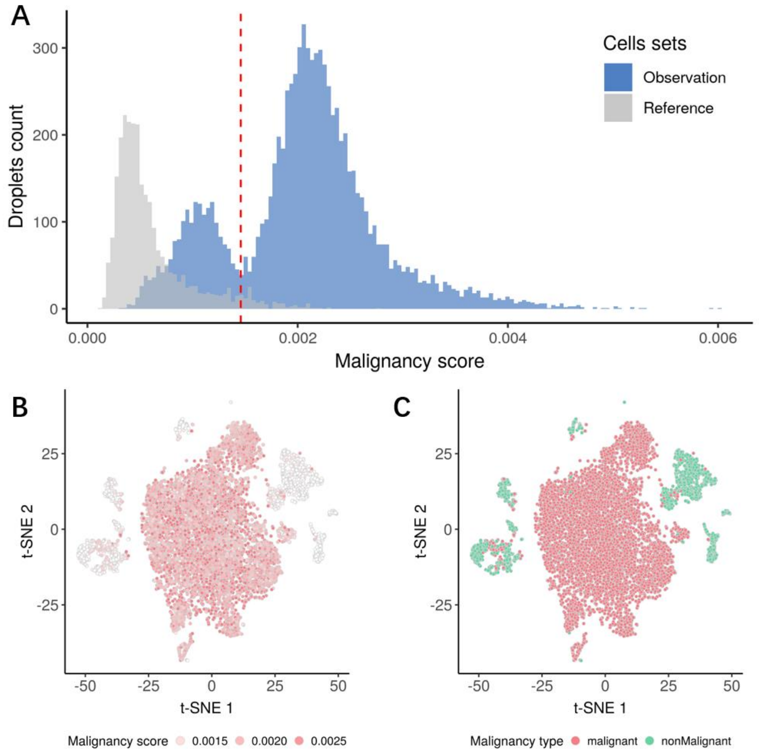


**Figure S6**. The visualization results of cell malignancy estimation in scCancer.

**(A)** The distribution of malignancy scores of observation (blue) of reference (grey) cells. The red dash line marks the identified threshold.

**(B)** A t-SNE plot shows the cells' malignancy score.

**(C)** A t-SNE plot shows the cells' malignancy type (red means malignant; green means non-malignant).

### S7. Tumor cell selection

In order to perform following intra-tumor heterogeneity analyses, we selected tumor cell clusters based on the results of cell type annotation (section S5) and cell malignancy estimation (section S6). We thought a cluster as tumor cells if its epithelial cell percent was larger than 60% and its malignant cell percent was larger than 90%.

### S8. Cell cycle analysis

To estimate cell cycle, we took advantage of a list of G2/M and S phase gene makers, and defined their relative average expression as cell cycle score. This step was implemented with the function "AddModuleScore" in Seurat [3].

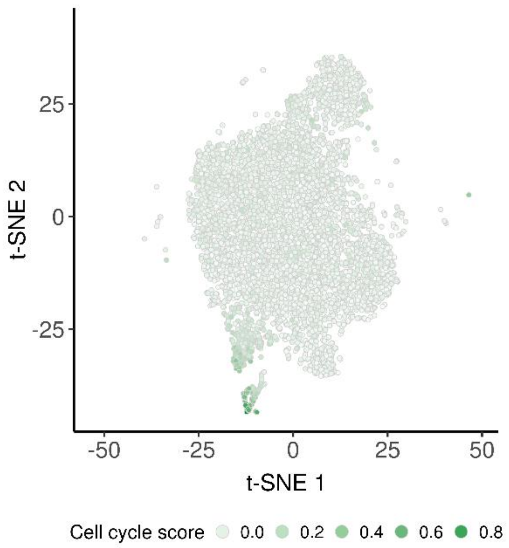


**Figure S7.** A t-SNE plot shows the cell cycle scores of the identified tumor clusters cells.

### S9. Cell stemness analysis

To analyze intra-tumor stemness heterogeneity, we referred to the workflow of Malta et al and estimated a stemness score for each cell [7]. In particular, we collected a stem/progenitor cells RNA-seq data from progenitor cell biology consortium (PCBC) dataset of Synapse (https://www.synapse.org/#!Synapse:syn1773109/wiki/54962), which contained an expression matrix with 12952 genes by 78 stem cell samples. Then, we trained a one-class logistics regression model (OCLR) on it using R package gelnet [5]. As a result, we got a stemness signature vector, which recorded the weights of 12952 genes for stemness. Using this signature, we calculated Spearman correlation with single cells' expression profiles, and defined the min-max normalized coefficients as the final stemness scores. The mathematical description of OCLR is similar to our cancer microenvironment cell type annotation step in supplementary section S5. And the detail of the workflow to train stemness signature can be found in the introduction of Malta et al (http://tcgabiolinks.fmrp.usp.br/PanCanStem/mRNAsi.html).

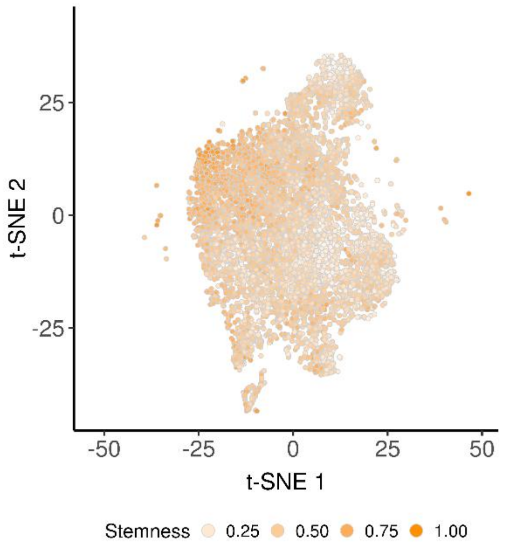


**Figure S8.** A t-SNE plot shows the stemness scores of the identified tumor clusters cells.

### S10. Gene set signature analysis

Analyzing in pathway or other gene set level may reveal more information on cells' expression characteristics. So, we also provided two optional methods to perform gene set signature analysis in scCancer. One is gene set variation analysis (GSVA) [8], where we applied GSVA function with its default setting on the scaled expression matrix of highly variable genes and calculated gene set scores for each cell. The other is calculating relative average expression levels as the scores, where we used the function "AddModuleScore" of Seurat and scaled the output matrix across gene sets by subtracting means and dividing standard deviations. By default, scCancer calculates signature scores of 50 hallmark gene sets from MSigDB (http://software.broadinstitute.org/gsea/msigdb/) and users can also input gene sets they are interested in.

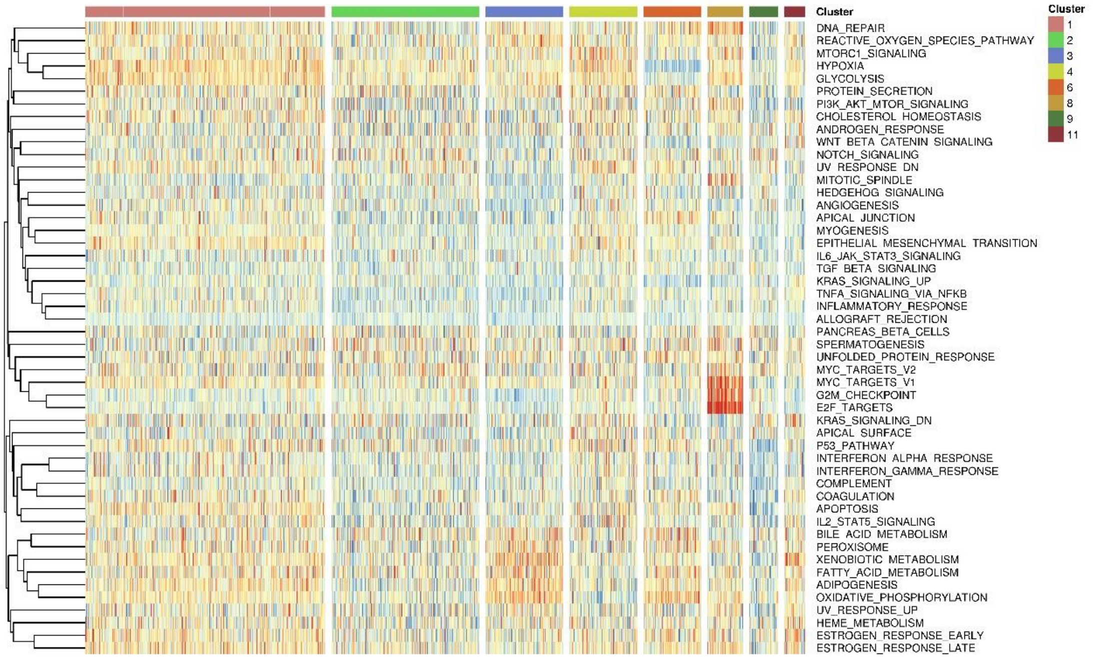


**Figure S9**. The visualization results of gene set signature analysis in scCancer. A heatmap shows the expression profiles of input gene sets (here is 50 hallmark gene sets). The color of top bar represents different clusters. And each row represents a gene set.

### S11. Expression programs identification

In order to identify potential expression programs by unsupervised ways, we also provided the function of non-negative matrix factorization (NMF). Firstly, we centered the log-transformed expression matrix by subtracting the means for each gene. Then we set the negative values of the matrix to zero. By applying NMF on it, we got two decomposed matrixes. The left one was gene by program, which we used to decide the related genes of programs. And the right one was program by cell, which was visualized by heatmap in scCancer to present cells' variable expression programs (Fig S10).

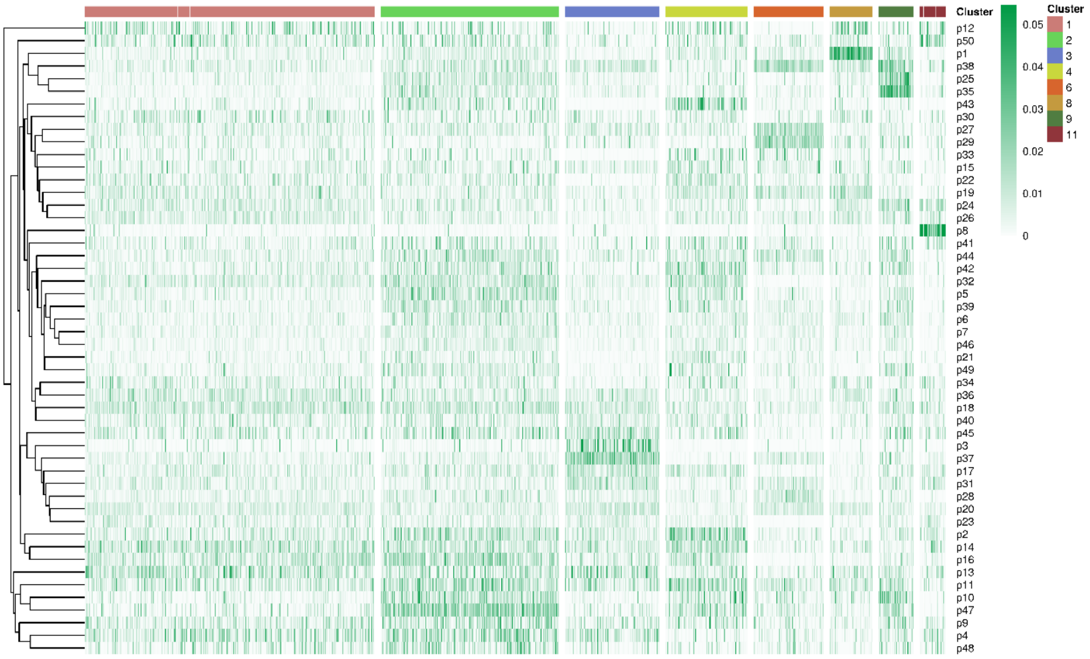

**Figure S10**. The visualization results of expression programs identification in scCancer. A heatmap shows the expression profiles of identified programs. The color of top bar represents different clusters. And each row represents a program.

### S12. Report generation

After all the steps, scCancer will save all the figures and results in a folder and generate three well-designed HTML reports from our R Markdown templates automatically. The first, named "report-cellRanger.html", is copied from the output of cell ranger. The second, named "report-scStat.html", records all results of statistics for cells and genes, which can be used to QC. The third, named "report-scAnno.html", records all the results of follow-up cell characteristics annotation. An example of the latter two reports can be found in the vignettes of scCancer.

### S13. The manual for scCancer

Our R package scCancer can be download and installed from GitHub (https://github.com/wguo-research/scCancer) and the detailed instruction is at the project wiki (https://github.com/wguo-research/scCancer/wiki/scCancer-vignettes).
